# A systematic analysis of the joint effects of ganglion cells, lagged LGN cells, and intercortical inhibition on spatiotemporal processing and direction selectivity

**DOI:** 10.1101/2024.01.31.578157

**Authors:** Reńe Larisch, Fred H. Hamker

## Abstract

Simple cells in the visual cortex process spatial as well as temporal information of the visual stream and enable the perception of motion information. Previous work suggests that the direction selectivity of V1 simple cells is associated with a temporal offset in the thalamocortical input stream through lagged and non-lagged cells of the lateral geniculate nucleus (LGN), but alternatively, it may also result from intercortical inhibition. While there exists a large corpus of models for spatiotemporal receptive fields, the majority of them built-in the spatiotemporal dynamics by utilizing a combination of spatial and temporal functions and thus, do not explain the emergence of spatiotemporal dynamics on basis of network dynamics emerging in the retina and the LGN. In order to better comprehend the emergence of spatiotemporal processing and direction selectivity, we used a spiking neural network to implement the visual pathway from the retina to the primary visual cortex. By varying different functional parts in our network, we demonstrate how the direction selectivity of simple cells emerges through the interplay between two components: tuned intercortical inhibition and a temporal offset in the feedforward path through lagged LGN cells. We observe that direction-selective simple cells are linked to a particular spiking pattern in a local excitatory-inhibitory circuit: If the stimulus moves in the non-preferred direction of a simple cell, inhibitory neurons with a different spatial position or tuning spike earlier, preventing the simple cell to spike. However, in the preferred direction, these inhibitory cells spike later, enabling the simple cell to spike.

## 1 Introduction

Simple cells in the primary visual cortex (V1) are selective for a wide range of features such as orientation, spatial frequency (Hubel & Wiesel, 1968), or stimulus movement direction (Hubel & Wiesel, 1959; Y. J. Kim et al., 2022; Wilson et al., 2018), characterized by their receptive field. The ON- and OFF-fields of a simple cell receptive field are typically not static but change on a millisecond time scale, known as a spatiotemporal receptive field (STRF) (McLean & Palmer, 1989; DeAngelis et al., 1995). The primary visual cortex is not the first stage showing such dynamic receptive fields, also thalamocortical cells in the lateral geniculate nucleus (LGN) (DeAngelis et al., 1995; Cai et al., 1997; Suematsu et al., 2013) and ganglion cells in the retina (Gollisch & Meister, 2010) reverse their receptive field polarity over time. Depending on how the spatial profile of the receptive field changes over time, a STRF can be classified as separable or inseparable (DeAngelis et al., 1993a). A separable STRF can be mathematically modeled by the multiplication of two independent functions, one for the space domain to describe the spatial selectivity of the cell, and one for the time domain to account for the switch in the polarity of the spatial receptive field (DeAngelis et al., 1995). In contrast, an inseparable STRF shows a transition in space over time (DeAngelis et al., 1993a).

To understand the emergence of spatiotemporal processing and direction selectivity in V1, different model approaches have been published, focusing on dynamics in early stages as well as intercortical circuit motifs. One of the first models accounting for separable and inseparable STRFs is the energy model from Adelson & Bergen (1985). While their model can reproduce different types of STRFs and detect stimulus motion, it does not consider biological mechanisms or circuit motifs.

A more biologically grounded approach utilizes the temporal offset between different cells in the LGN to explain the emergence of a variety of simple cell STRFs (Hamada et al., 1997; Wimbauer, Wenisch, Miller, & van Hemmen, 1997; Miyashita et al., 1997; La Cara et al., 2003). In these models, the spatial profile of the LGN cells is modeled either as a difference of Gaussian (Miyashita et al., 1997; La Cara et al., 2003; Chizhov & Merkulyeva, 2020; Chariker et al., 2021) or as a Gabor-function (Wimbauer, Wenisch, Miller, & van Hemmen, 1997), while the temporal function is bi-phasic to simulate STRFs of the LGN (Hamada et al., 1997; Wimbauer, Wenisch, Miller, & van Hemmen, 1997; Miyashita et al., 1997; La Cara et al., 2003). Adding a delay to the temporal function of some LGN cells leads to V1 simple cells with inseparable STRFs (Hamada et al., 1997; Wimbauer, Wenisch, Miller, & van Hemmen, 1997; Miyashita et al., 1997; La Cara et al., 2003). LGN cells with a delayed response are called lagged LGN cells and have been observed in cats (Saul & Humphrey, 1990) and monkeys (Saul, 2008b). Similar to the assumption of Adelson & Bergen (1985), that simple cells with an inseparable STRF are motion detectors, it has been demonstrated by models with this functional approach that simple cells with an inseparable STRF show a higher direction selectivity (Hamada et al., 1997; Wimbauer, Wenisch, Miller, & van Hemmen, 1997), suggesting that the temporal offset in the LGN input leads to direction selectivity or at least provides a baseline selectivity which is sharpened through intercortical inhibition (Miyashita et al., 1997; La Cara et al., 2003; Ursino et al., 2007; Okajima, 2014; Chizhov & Merkulyeva, 2020). Other model studies demonstrated that this required temporal offset can be caused by the interaction between LGN cells with a sustained response profile and cells with a transient response profile (Chizhov & Merkulyeva, 2020), or with a delay between ON- and OFF-LGN cells (Chariker et al., 2021). While these model studies are successful in modeling simple cell spatiotemporal receptive fields and direction selectivity, the temporal dynamics of LGN cells are implemented with a temporal function without incorporating explicit local neuronal dynamics.

Another class of models assumes that direction selectivity is caused alone by intercortical inhibition instead of a temporal offset in the LGN responses (Wenisch et al., 2005; Freeman, 2021). Wenisch et al. (2005) demonstrated in a spiking neural network, consisting of excitatory and inhibitory neurons to mimic the primary visual cortex, how the dynamic between lateral excitatory and inhibitory currents leads to direction selectivity. They use an asymmetric spike-timing-dependent plasticity rule to demonstrate how the lateral excitatory synapse develops a spatial asymmetry causing direction selectivity (Wenisch et al., 2005). While the simple cell selectivity in Wenisch et al. (2005) was predefined and only the lateral excitatory synapses are plastic, Freeman (2021) used a more detailed rate-based model from the photoreceptors in the retina up to the primary visual cortex. With a Hebbian-like learning scheme, they tuned the selectivity of excitatory and inhibitory cortical cells to learn orientation preference maps. The value of inhibitory synapses, however, increases linearly during the learning phase, independent of pre-and postsynaptic activity. Inhibitory cells are connected to excitatory cells in their spatial neighborhood, weighted with a Gaussian function. Due to this, direction-selective excitatory cells show up at the edge between two subareas of homogeneous orientation preference. Receiving inhibition from inhibitory neurons with a similar and different orientation selectivity in the neighborhood leads to a temporal offset between the arriving inhibitory currents at the preferred and non-preferred direction (Freeman, 2021). While cat visual cortex is indeed organized in orientation-dependent functional maps (Ohki et al., 2006), other species with direction-selective cells, e.g. rats, do not have such maps, so inhibitory cells in the neighborhood of a direction-selective cell do not show a temporal bias due to orientation preference (Ohki et al., 2005). While models of Wenisch et al. (2005) and Freeman (2021) show the importance of intercortical inhibition for direction selectivity, both consider downstream areas (such as the retina and the LGN) as simple relays of spatial information, ignoring their temporal dynamics. Further, a few simplifications may affect the modeled conclusions, such that the synapses from the inhibitory to the excitatory cells are initialized randomly (Wenisch et al., 2005) or their weight increases linearly during the learning phase, without taking the pre-and post-synaptic activity into account (Freeman, 2021). Concluding, their studies do not proof that thalamocortical temporal dynamics compared to tuned feedback inhibition are irrelevant for the emergence of simple cell direction selectivity.

While most models focus on a single cause to explain the occurrence of direction selectivity, the variety of possible explanations suggests rather multiple, presumably redundant mechanisms. To improve our understanding of the emergence of STRFs and to identify necessary network dynamics of direction selectivity in V1, we performed a systematic analysis of different network motifs in a spiking neural network, modeling the visual pathway from the retinal ganglion cells to the primary visual cortex, considering different temporal dynamics along the pathway. Our model is inspired by the visual pathway of primates, but when appropriate, we refer to findings in other species. We assume that a delay in the surround field of the ganglion cells leads to a temporal decoupling of the processing of visual information along the center and the surround, sending a phasic signal to the cortex. To avoid a bias in the emergent direction selectivity due to temporal dynamics in the ganglion cells we did not implement direction selectivity in the retina. The connectivity matrix from the LGN cells to the V1 simple cells and inhibitory interneurons has been obtained by prior training on static natural scenes to ensure a realistic connectome in our network.

While several experimental studies have reported direction-selective ganglion cells in the retina of different mammals (Weng et al., 2005; Hillier et al., 2017; Y. J. Kim et al., 2022) and in LGN (Xu et al., 2002; Piscopo et al., 2013; Stacy et al., 2023), the rather low number of direction-selective cells in retina (Y. J. Kim et al., 2022) and in LGN (Xu et al., 2002) suggests a weak influence on the direction selectivity of cortical cells. Additionally, it has been shown that disrupting direction selectivity in the retina did not influence the amount of direction-selective cells in V1 but removed a bias towards the four cardinal directions (Hillier et al., 2017). Due to this, we assume the ganglion cell direction selectivity plays only a minor role in simple cell direction selectivity, and thus, we did not consider them in our neural model. Through systematic modifications, we investigate how the tuning of intercortical inhibition and the temporal offset caused by non-lagged and lagged LGN cells influence the emergence of direction selectivity. Our results suggest that tuned intercortical inhibition alone enables a sharp orientation tuning but shows a symmetric, bidirectional response profile. A temporal offset in the thalamocortical input, as introduced between lagged and non-lagged LGN cells, is necessary to break this symmetry and enable the emergence of unidirectional selectivity. Further, resulting network dynamics are consistent with observations, such as that pyramidal cells with high unidirectional selectivity receive inhibition later than their excitatory component when stimulated with a stimulus moving in the preferred direction, while when the stimulus moves in the non-preferred direction, the excitatory and inhibitory currents arrive at the same time and cancel each other out (Priebe & Ferster, 2005; Li et al., 2014).

To the best of our knowledge, we present the first spiking neural network utilizing a delay in the surround fields of retinal ganglion cells (RGC), lagged LGN cells, as well as intercortical inhibition to enable spatiotemporal processing of V1 simple cells. By performing a systematic analysis, we demonstrate that direction selectivity can emerge through the combination of lagged LGN cells and tuned intercortical inhibition.

## 2 Methods

The spiking neural network has been implemented in Python 3.8 and by the ANNarchy simulator (v4.7.1) (Vitay et al., 2015). Source code of the spiking network and evaluations are available at https://github.com/hamkerlab/Larisch2024 Spatiotemporal Processing.

### 2.1 Network architecture

The model is based on the V1 layer 4 model proposed in our previous work (Larisch et al., 2021), but extended by input processing to enable the emergence of spatiotemporal behavior. Due to this, the network implements three different stages of the visual stream, from the retina via the lateral geniculate nucleus to layer 4 of the primary visual cortex (see **Fig. 1**). Although the architecture of our network is inspired by the visual pathway of primates, we did not implement a detailed species-specific model and rather investigate the conceptual reasons for spatiotemporal behavior.

**Figure 1:**
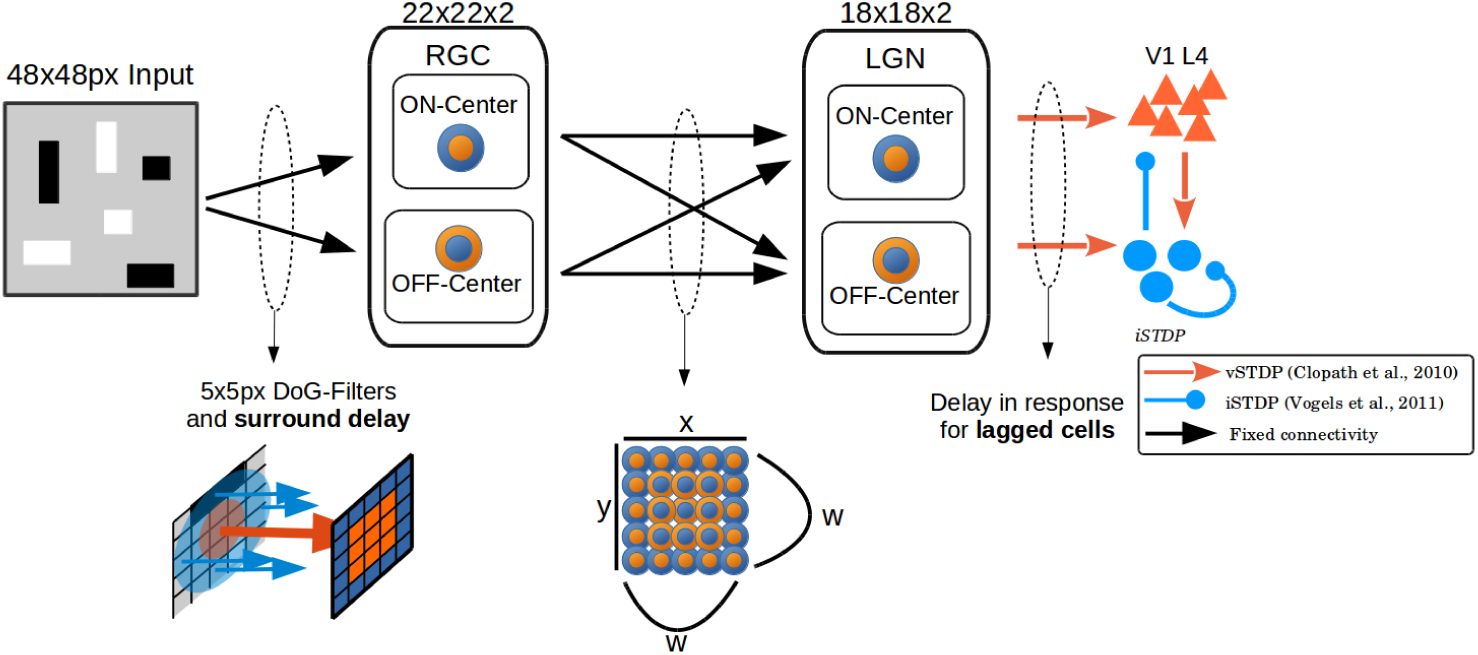
Network architecture. A 48*×*48 pixel input is presented to the input layer and sent to two populations of retinal ganglion cells (RGC). One population consists of ON-center cells and the other of OFF-center cells to mimic the ON- and OFF-path. The input weights of the RGCs are modeled as a Difference of Gaussian so that each RGC receives a 5*×*5 pixel input. A delay can be added to the weights of the DoG surround field. A LGN cell receives input from a grid of 5*×*5 RGC to preserve the spatial arrangement. To enable the center-surround antagonism in LGN, an LGN cell receives input from 3*×*3 ganglion cell grid of the same polarity for the center, and a ring of 16 ganglion cells with the opposite polarity for the surround. The weights from the ganglion to the LGN cell follow a spatial Gaussian distribution, with higher weights in the center and lower weights in the surround field. Both LGN populations are connected to a population of excitatory and inhibitory cells, whose connections are learned on natural scenes. A lag in the transfer of an LGN spike can be added to convert LGN cells to lagged LGN cells.

**Figure 2:**
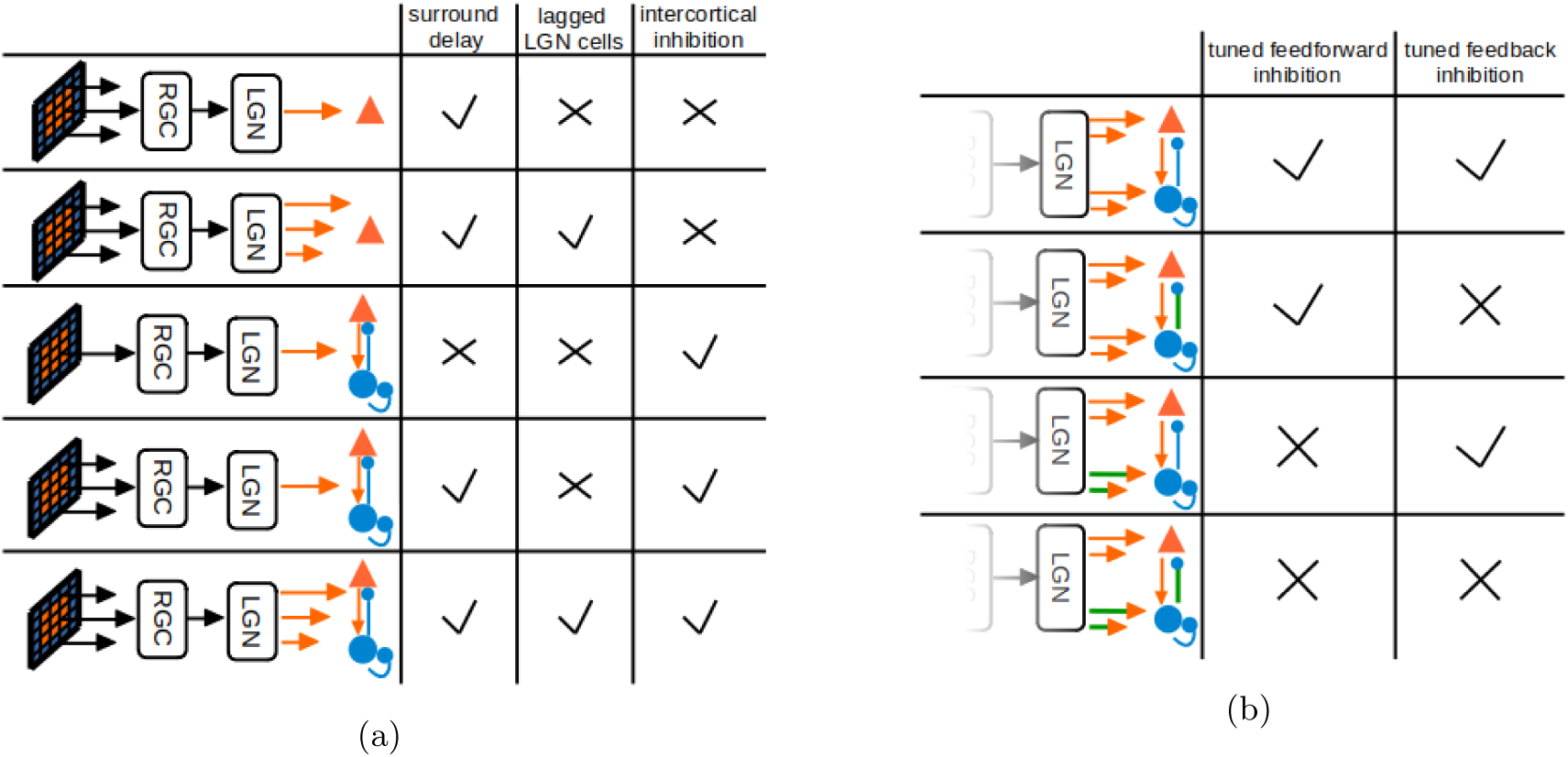
Pictograms of network modifications to evaluate their influence on direction selectivity. (a) Network modifications to analyze the influence on the emergence of STRFs and direction selectivity. We varied the delay in the surround field of the RGC receptive fields, adding lagged cells to the LGN population and adding tuned intercortical inhibition. (b) Network modifications to analyze the influence of tuned feedforward and feedback inhibition on direction selectivity. While Orange arrows indicating tuned excitatory and blue arrows tuned inhibitory synapses, green arrows indicating connections that are shuffled to remove the structure of the specific connection. Staggered arrows indicating a delay in either the synapses or the neuron responses.

#### 2.1.1 Retina

The first stage implemented in the network contains the retinal ganglion cells (RGC). We model the input weights for a RGC with a two-dimensional difference of Gaussian (DoG) function, a common function to model the spatial center-surround organization of the RGC receptive fields (Enroth-Cugell et al., 1983; Troy & Shou, 2002), where the ON-fields provide excitatory input and the OFF-fields inhibitory input. This mimics the joint effect of the bipolar cells and the inhibitory horizontal cells, and the bipolar cells and inhibitory amacrine cells in the retina (Diamond, 2017; Baden et al., 2020). The receptive fields sample from a 5*×*5 pixel input. With a spacing of 2 pixels between two neighboring cells, the complete population receives a monochrome 48*×*48 pixel input. The DoG functions of two adjacent cells have an overlap of 3 pixels. The model implements the ON- and OFF-pathway via two populations of RGCs. The ON-pathway consists of ON-centered cells and the OFF-pathway of OFF-centered cells. Thus, the complete RGC layer consists of 22*×*22*×*2 cells arranged spatially on a 2D-grid to process the input. The RGCs are modeled with adaptive integrate-and-fire neurons (Brette & Gerstner, 2005), receiving a continuous excitatory (via the ON-field) or inhibitory (via the OFF-field) input current.

We add a delay to surround fields of the RGCs, as reported for bipolar and ganglion cells in the retina of different mammalians (Kilavik et al., 2003; Tokutake & Freed, 2008). Its value was drawn from a discrete uniform distribution between 30 ms and 50 ms, relative to the center weights. While the reported surround delay in the primate retina is shorter (6 ms to 8 ms) (Benardete & Kaplan, 1997; Kilavik et al., 2003), our chosen range is similar to the range used by Y. J. Kim et al. (2022) to fit the center-surround dynamic of primate bipolar cells, which provide the feedforward input to ganglion cells.

#### 2.1.2 Lateral geniculate nucleus

The LGN layer consists of two LGN populations, implementing the ON- and OFF-pathway, each with 18*×*18 LGN cells, arranged on a 2D grid. To realize the center-surround antagonism, each LGN cell receives input from the RGC population with the same polarity for the center field and from RGC cells with an opposite polarity for the surround field, similar as it is reported for retinogeniculate connections in cats (Suematsu et al., 2013). For example, an ON-center with OFF-surround LGN cell is connected to 9 (in a 3*×*3 grid) RGCs in the ON-population and to 16 (to assemble the surround field) cells in the OFF-population (and vice versa for an OFF-centered, ON-surround LGN cell).

As studies reported that some LGN cells respond with a delay of 30 ms to 70 ms to the stimulus onset (so-called lagged LGN cells) (Saul & Humphrey, 1990; Saul, 2008a; Vigeland et al., 2013), we implemented a synaptic delay on the outgoing synapses of a subgroup of LGN neurons to create a lag in the transmission of the LGN spikes, accounting for observed lagged LGN cells.

#### 2.1.3 Layer 4 of primary visual cortex

Both LGN populations (ON- and OFF-center) project to a population of excitatory and inhibitory neurons resembling layer 4 of the primary visual cortex (V1). The excitatory population consists of 324 neurons and the inhibitory one of 81 neurons to match the 4:1 ratio, as reported for different species as in the striate cortex of cats (Gabbott & Somogyi, 1986), monkeys (Beaulieu et al., 1992) and in rodents (Markram et al., 2004). Inhibitory neurons receive additional input from the excitatory neurons, sending inhibitory input back to the excitatory neurons, and they are connected to other inhibitory neurons of their population (no self-inhibition). To ensure the emergence of Gabor-like receptive fields in the excitatory and inhibitory population, and of a structured connectivity between these two populations we pre-trained the weights to and between both populations in a separate training step. In this training step, 400,000 randomly chosen patches of natural scenes (Olshausen & Field, 1996) are presented. The learning of these connections was done separately, without the RGC population, and with a Poisson neuron model in the LGN. A detailed description of the training and properties of the resulting network can be found in Larisch et al. (2021). After that, no further learning took place and every other connection in this network is fixed. As the synapses of the network were initialized randomly for the pre-training, we trained 5 different network instances to ensure, that the observed dynamics were not caused by random effects through the initialized synapses. Each network instance represents one complete model with a different set of pre-trained weights. We performed all analyses on each of the five network instances.

#### 2.1.4 Neuron model

All neurons in the RGC and LGN are implemented on the basis of the adaptive exponential integrate- and-fire neuron model (AdEx) as described in Brette & Gerstner (2005). The membrane potential (*V* ) develops as described in Eq.1 and depends on the input *I*, representing the sum over the excitatory and inhibitory input current into the neuron, and the adaptive current *w* (see Eq.2). If *V* exceeds the spiking threshold (*V_T_* ) the exponential term increases promptly resulting in a rapid increase of the membrane potential. After each spike, *w* is increased by a value *b*, and *V* is set to *V_r_*.

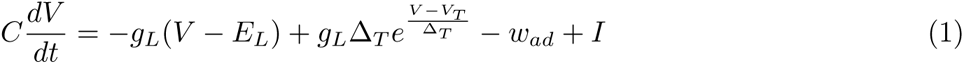

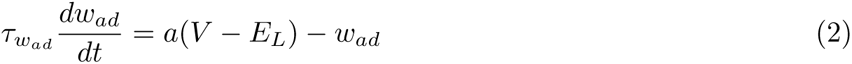

Additional fixed parameters in the AdEx model are *C* as the membrane capacitance, *g_L_*as the leak conductance, *E_L_*as the resting potential, and Δ*_T_* as the slope factor. The parameter values are taken from Naud et al. (2008) (see Tab. 1 for parameter values).

**Table 1:**
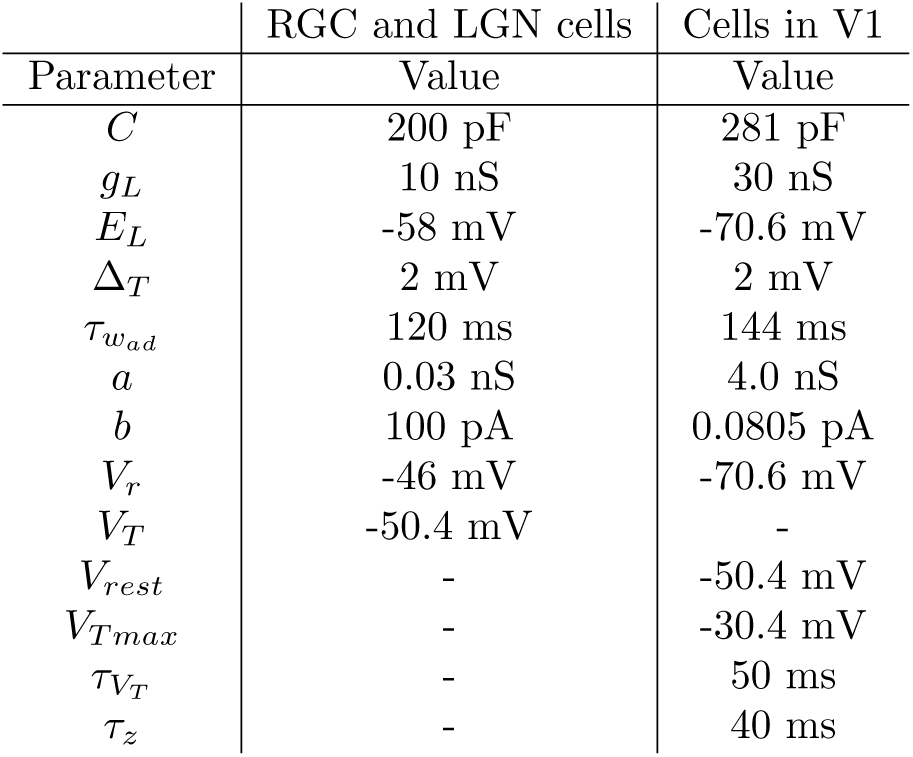
Parameter for the AdEx neuron model in the LGN and RGC population shown in the left column. Values are chosen to implement a bursting behavior (Naud et al., 2008). Right column shows parameter for the AdEx neuron model in the excitatory and inhibitory population.

Due to the chosen learning rules in the separate learning phase of the layer 4 model part, the excitatory and inhibitory population consists of AdEx neurons following the implementation in Clopath et al. (2010) (see Tab.1 for parameter values). In this slightly different neuron model, the membrane potential is extended with a depolarizing afterpotential (*z*), which increases after each spike by *I_sp_* and decays with *τ_z_*to zero (Eq.3). Additionally, the fixed spiking threshold is replaced by an adaptive threshold, which is set after each spike to *V_T max_* and decays with a time constant of *τ_V__T_* back to the lower value of *V_rest_*(Eq.4).

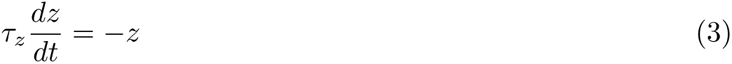

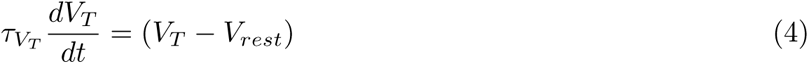

### 2.2 Network modifications

To investigate how the circuit dynamics are influenced by intercortical inhibition or the temporal offset between lagged and non-lagged LGN cells, we measured the simple cell direction selectivity on five different modifications of the network, modifying the delay in the feedforward synapses of the RGC, the LGN responses and the intercortical inhibition (**Fig.2a**).

To study how the network dynamics influence the emergence of the STRFs, we modified the delay in the surround field of the retinal ganglion cells. Further, we add a delay of 50 ms to the response of 50% of the LGN cells to implement a temporal offset in the thalamocortical input to the simple cells.

The influence of the temporal offset and inhibition on the direction selectivity has been investigated by the following modifications: In the first modification, all LGN cells respond at the same time (so no lagged LGN cells exist) and intercortical inhibition is not present in the network. For the second modification we added lagged LGN cells to implement a temporal offset in the thalamocortical input stream, still without intercortical inhibition. Finally, we added tuned intercortical inhibition to the network and measured direction selectivity once with and once without lagged LGN cells.

To evaluate the influence of tuned intercortical inhibition on direction selectivity, we compare the network with tuned feedforward and feedback inhibition with three variants in which in some connections the tuning has been removed by shuffling the weights (**Fig.2b**). For the first variant, we removed the tuning in the feedback synapses from the inhibitory to the excitatory population. In the second variant, we removed the tuning in all excitatory synapses to the inhibitory population (from LGN as well as from the excitatory population), removing the spatial tuning of inhibitory neurons. Finally, we removed the tuning of all synapses to the inhibitory population and in the inhibitory feedback synapses. We did not untune the lateral inhibitory synapses within the inhibitory population, as previous work demonstrated that this will not have a strong effect on the simple cell responses (Larisch et al., 2021).

### 2.3 Analysis methods

#### 2.3.1 Spatiotemporal receptive fields

An established method for measuring the spatiotemporal receptive fields is the reverse correlation technique (McLean & Palmer, 1989; DeAngelis et al., 1993a, 1995; Ohzawa et al., 1996). We presented 200,000 randomly chosen bright or dark bars and recorded all time points where a cell spiked during the presentation. Each bar was presented for 50 ms, its position was chosen randomly and the size varied between one and eight pixels in height and width. To obtain a 3D-response profile, we summed over the stimuli presented in the last 250 ms before a spike occurred and divided it by the number of spikes.

#### 2.3.2 Quantify the inseparability of STRFs

To measure if the STRF of an excitatory cell is either separable or inseparable, we first calculated the frequency spectrum of each neuron via a Fourier transformation (DeAngelis et al., 1993a). The amplitudes for positive and negative temporal frequencies should be equal for separable STRFs, and different for inseparable STRFs (DeAngelis et al., 1993a; Ohzawa et al., 1996; T. Kim & Freeman, 2016). Thus, the difference between the temporal frequency amplitudes (TFD) quantifies the ”inseparability” of the excitatory cells (see **Eq.5**), with *A_p_*as the maximum amplitude value of positive frequency and *A_n_* maximum amplitude of the negative frequencies. For separable STRF the TFD value is zero, whereas a value closer to one indicates an inseparable STRF.

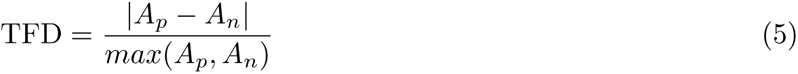

#### 2.3.3 Direction selectivity index

Direction selectivity is determined by using moving sinusoidal gratings (Peterson et al., 2004; Priebe & Ferster, 2005). To find the optimal moving sinus grating for each cell, we varied the spatial frequency from 1 *cycles/image* up to 5 *cycles/image* and the temporal frequency from 2 *cycles/second* up to 6 *cycles/second*. Each grating was presented for 20 different movement directions and moved for 3000 ms in each direction. We also determined the preferred orientation of each cell with a static, non-moving sinus grating. The preferred orientation together with the preferred spatial and temporal frequency is used to determine the two directions where the cell shows its highest response (preferred direction) and its lowest response (null or non-preferred direction). We calculate the direction selectivity index (DSI) as proposed by Peterson et al. (2004) (Eq.6):

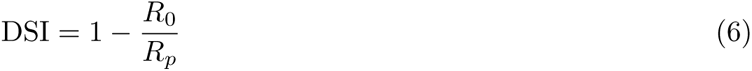

*R_p_*is the neuron activity (number of spikes) in the direction with the highest activity (preferred direction) and *R*_0_ is the neuron activity in the null (opposite) direction. Cells that are highly selective for one direction have a DSI closer to 1, and cells that a less selective for one direction have a DSI close to 0.

#### 2.3.4 Temporal offset between excitation and inhibition

To calculate the temporal offset (Δ*_T_* ) between the excitatory and inhibitory input currents we calculated the Pearson correlation between the excitatory input current and a time-shifted version of the inhibitory input current, similar to the method described in Okun & Lampl (2008). Therefore, the inhibitory current is shifted in time from -20 ms up to + 20 ms. The time shift where the correlation value is highest is used as Δ*_T_* to determine the arrival of inhibition relative to the arrival of excitation. A negative value of Δ*_T_* indicates a delayed arrival of the original inhibitory current, whereas a positive value indicates that the inhibitory input arrives earlier than the excitatory one.

To relate the temporal dynamics of the inhibitory cells to the similarity between their receptive field and those of excitatory cells, we first calculated the receptive field similarity between all excitatory and inhibitory cells (see below). Second, for each excitatory cell, we binned inhibitory neurons with respect to their similarity to five bins. Third, we calculated the peristimulus time histogram (with a time bin of 2 ms and over 5 stimulus repetitions) for each inhibitory neuron in one bin and multiplied it by the weight value of the inhibitory feedback synapse. By averaging over all inhibitory cells in each bin, we calculate the expected inhibition an excitatory cell receives over the time of a stimulus presentation. Finally, we averaged the excitatory input current over 2 ms time bins and calculated the temporal offset as described above.

#### 2.3.5 Selectivity similarity

To measure if a pair of excitatory and inhibitory cells are similar in their selectivity, we used the feedforward weights (excitatory input synapses from the LGN) and calculated the cosine between them (see **Eq.7**), with *W_i_* as the feedforward weight vector of the excitatory neuron *i* and *W_j_* as the feedforward weight vector of the inhibitory neuron *j*.

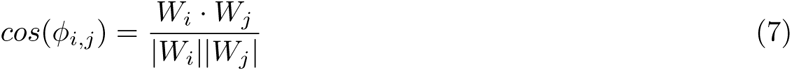

## 3 Results

### 3.1 Surround to center delay leads to spatiotemporal receptive fields

Our results show that the structure of a spatiotemporal receptive field (STRF) depends on the surround delay of the ganglion cell receptive field and the temporal offset between lagged and non-lagged LGN cells (**Fig. 3**). Adding a delay of 30 ms to 50 ms to the surround field of the ganglion cells leads to a polarity flip of the simple cell receptive fields along the time axis in the STRF, typical for separable STRFs (DeAngelis et al., 1993a) (**Fig. 4**). An additional lag of 50 ms in the responses of 50% randomly chosen LGN cells leads to the emergence of an inseparable STRF profile (**Fig. 3**).

**Figure 3:**
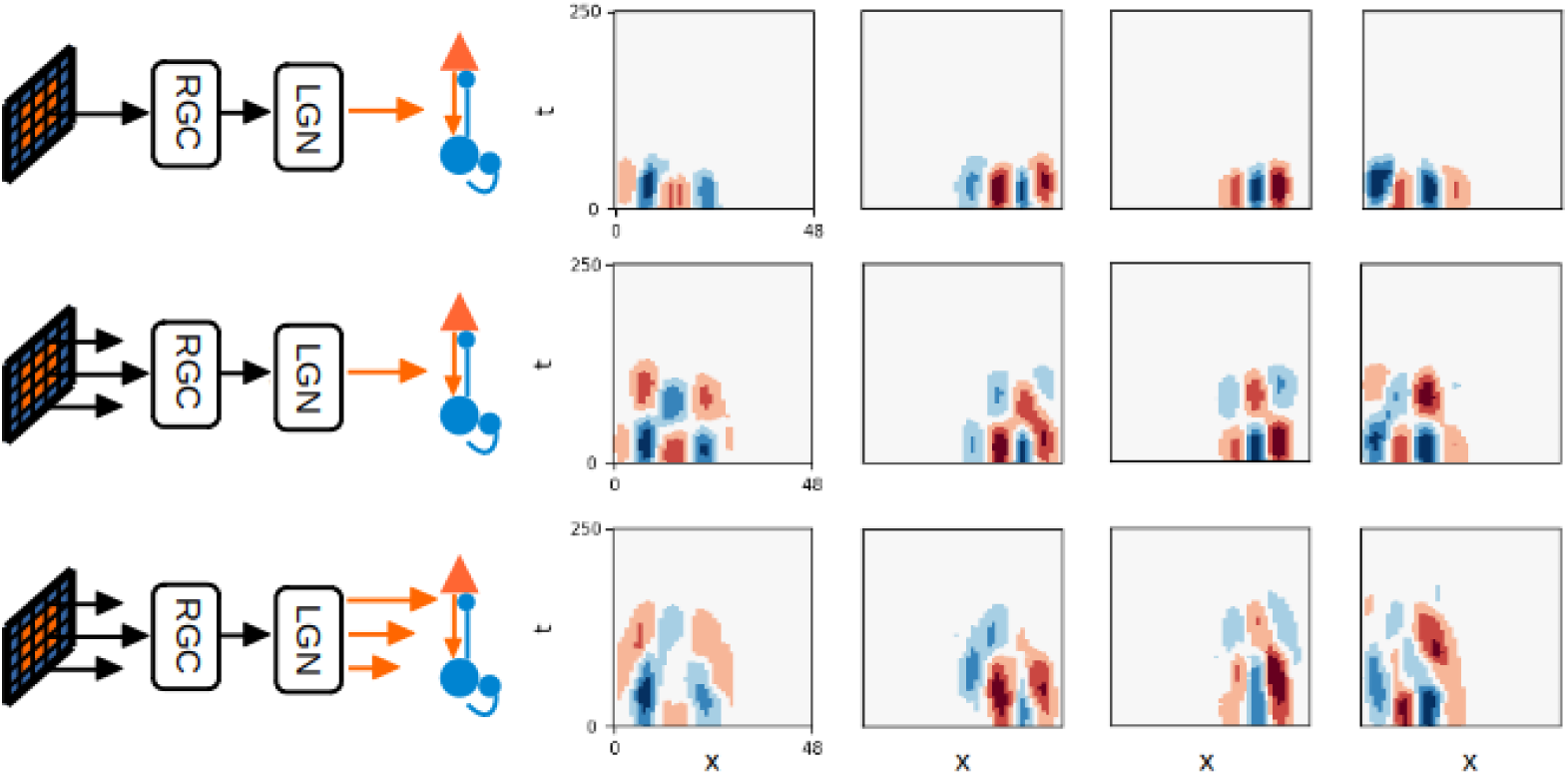
Spatiotemporal receptive fields. Each column shows the spatiotemporal receptive field (STRF) of one simple cell with different network configurations. If there is either a delay in the RGC surround field or a delay in the response of the LGN cells, there is no typical dynamic in the STRF (first row). With a delay in the RGC surround fields, separable STRFs emerge (second row). With a delay in the RGC surround fields and a lagged response for 50% of the LGN cells, the STRFs become more inseparable (third row).

**Figure 4:**
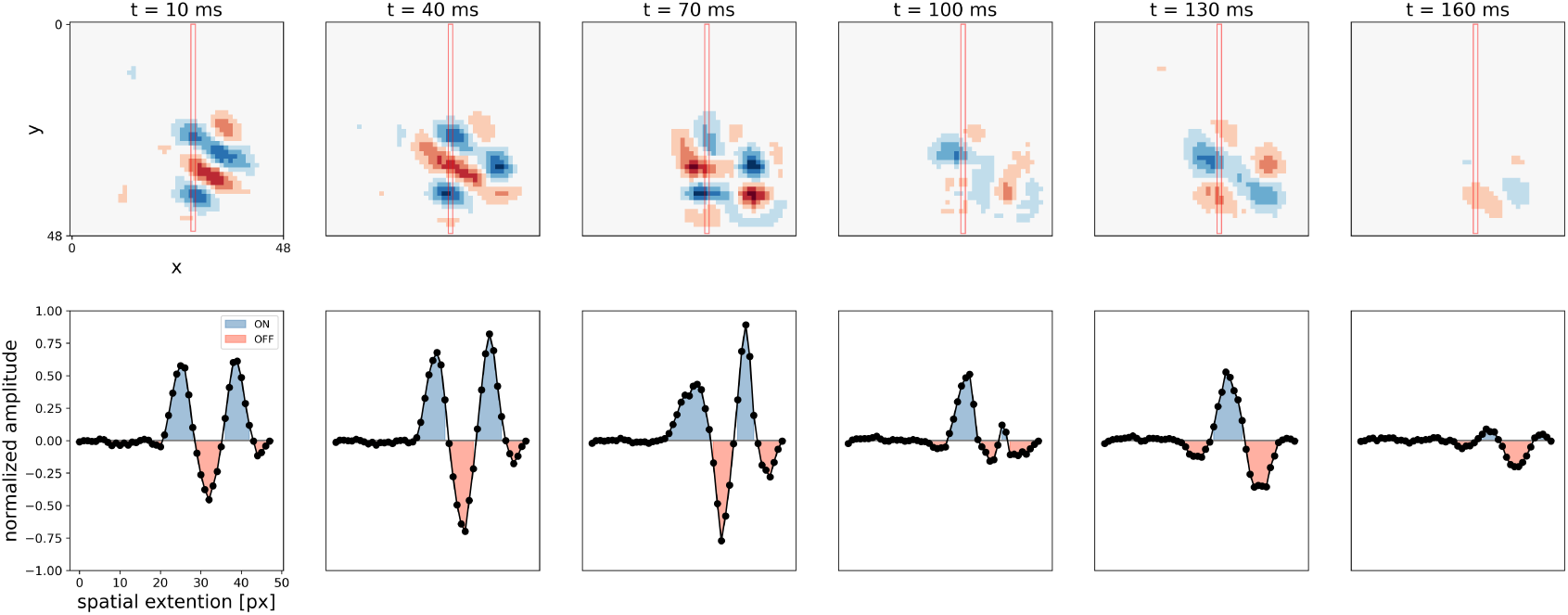
Temporal change of simple cells receptive fields. Temporal dynamic is indicated the changing polarities of the spatial receptive field between time point *t* = 10 ms and *t* = 130 ms. **Top:** Spatial receptive field of a randomly selected neuron, at different time steps, taken from the three-dimensional spatiotemporal response profile (see **Sec. 2.3.1**). **Bottom**: One-dimensional normalized response profile of the spatial receptive field for the corresponding time steps. The red rectangle in the image above indicates the position of the profile.

Experimental data suggests, that the frequency spectrum of a separable and inseparable STRF differ in terms of the amplitudes for the negative and positive temporal frequencies (DeAngelis et al., 1993a; Ohzawa et al., 1996; T. Kim & Freeman, 2016). For cells with a separable STRF, the amplitudes for positive and negative temporal frequencies are equal, whereas the amplitudes for an inseparable STRF differ.

We calculated the frequency spectrum of all STRFs in the model with and without lagged LGN cells, respectively, and observed that the amplitudes are equal for a separable STRF (**Fig.5a**) and differ for a more inseparable STRF (**Fig.5b**). To quantify the ”inseparability” of each STRF, we calculated the difference between the amplitudes of temporal frequencies (TFD). If the TFD value is closer to zero, it indicates a rather separable STRF and a value closer to one indicates a rather inseparable STRF (see **Eq. 5**). The TFD allows us to measure the effect of lagged LGN cells on the resulting distribution of separable and inseparable cells. We observe that without lagged LGN cells, most of the neurons have a low TFD, indicating that most neurons have separable STRFs (see **Fig.6a**). In contrast, lagged LGN cells lead to higher TFD values and so to more neurons with inseparable STRFs (see **Fig.6b**). Still, in either case we observe a range of TFD values, indicating a spectrum of different STRFs from totally separable to clearly inseparable ones.

**Figure 5:**
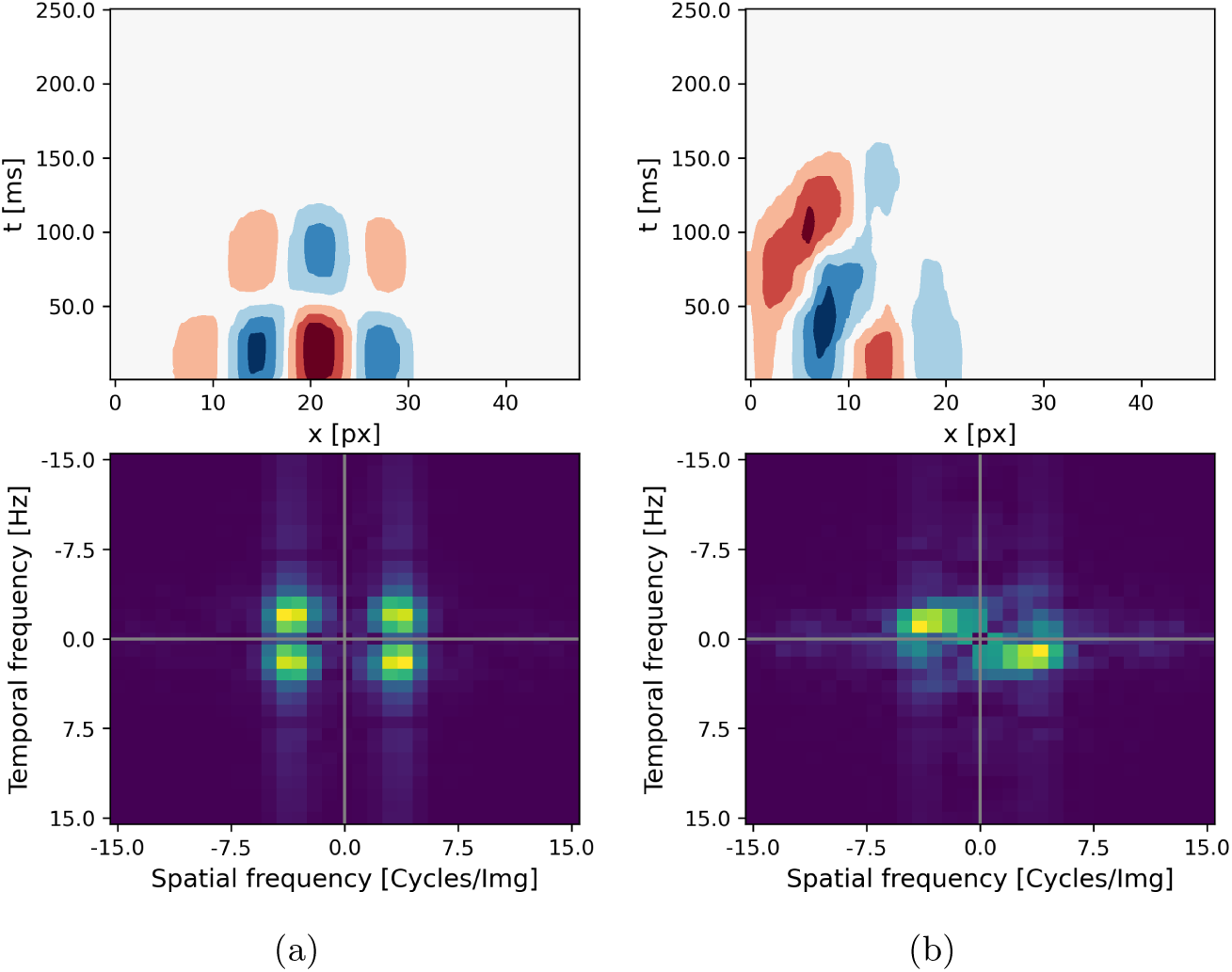
Inseparable STRF show different amplitudes for positive and negative temporal frequencies. (a) Separable spatiotemporal receptive field of one exemplary cell (top) and the corresponding frequency spectrum (bottom). **(b)** Inseparable spatiotemporal receptive field of one exemplary cell (top) and the corresponding frequency spectrum (bottom). The frequency spectrum of the cell with the inseparable spatiotemporal receptive field is only diagonal-symmetrical at the centered zero point.

**Figure 6:**
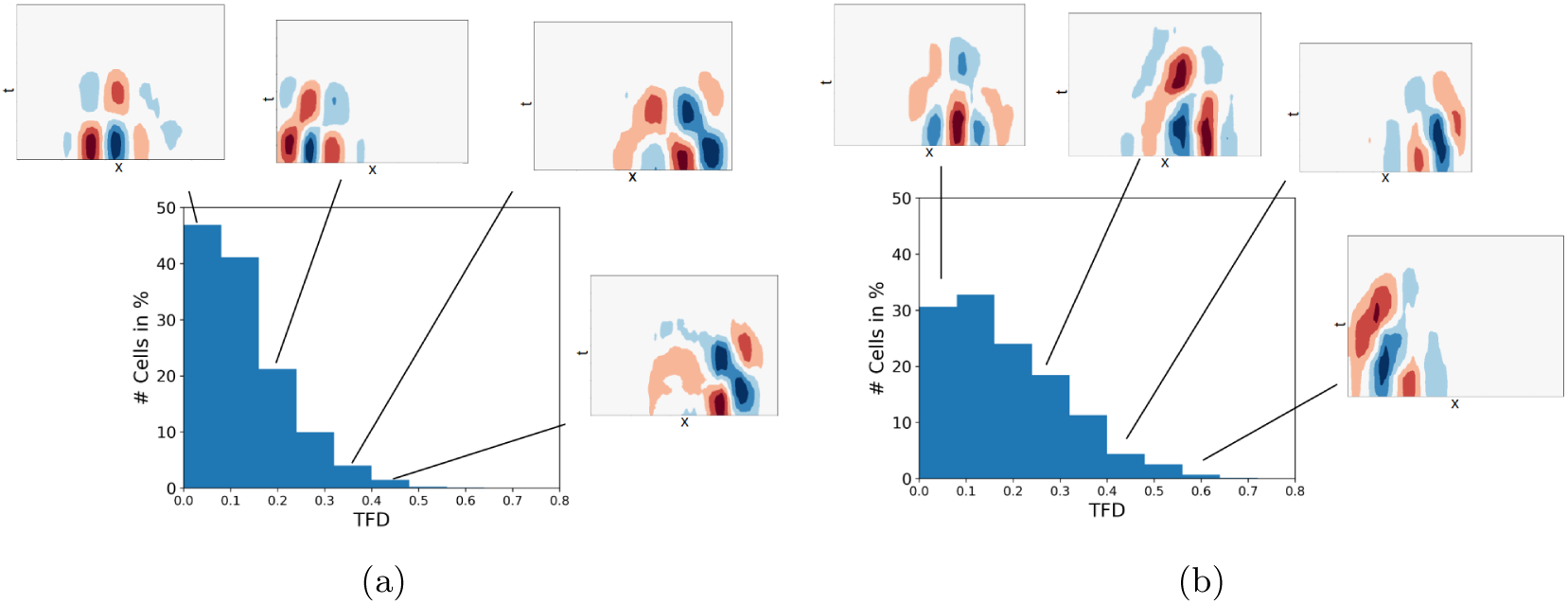
A more inseparable STRFs leads to a higher difference in the temporal frequency amplitudes (TFD). (a) Without any lagged LGN cells, most of the STRFs have a small TFD value and rather separable STRFs. **(b)** If 50% of the cells in the LGN population are lagged cells with a delay of 50*ms*, many simple cells tend to have inseparable STRFs indicated by a higher TFD value.

### 3.2 Direction selectivity through lagged LGN cells and intercortical inhibition

We have seen an influence of lagged LGN cells on the inseparability of the STRF in our model. It has been suggested by previous model studies, that inseparable STRFs are more direction selective, linking lagged LGN directly to the emergence of direction selectivity (Miyashita et al., 1997; Wimbauer, Wenisch, Miller, & van Hemmen, 1997). Furthermore, it has been suggested that intercortical inhibition could also cause direction selectivity (Priebe & Ferster, 2005; Freeman, 2021). To analyze in our model the influence of both, lagged LGN cells and inhibition, we measured the direction selectivity of the excitatory neurons under varying model configurations.

We varied the model by removing the temporal lag in the response of the lagged LGN cells or by deactivating the intercortical inhibition but leaving the delay in the ganglion surround fields intact (**Fig.7**). Without either lagged LGN cell or inhibition, we obverse that most of the simple cells show no or only weak direction selectivity. With 50% of the LGN populations being lagged cells (with a lagged response of 50 ms), but without inhibition, the DSI distribution does not vary. If the network contains no lagged LGN cells but active intercortical inhibition, more cells show a higher direction selectivity, leading to a more long-tailed distribution of the DSI. If both exist, lagged LGN cells and inhibition, even more neurons gain a higher direction selectivity, approaching a uniform distribution. These results suggest, that both, lagged LGN cells and inhibition, influence the emergence of direction selectivity. To analyze how both work together, we perform a qualitative evaluation by studying the response profile of four cells with different direction selectivity in the four different model configurations (see **Fig. 8**). The resulting response profiles show, that with blocked intercortical inhibition a broad tuning is observable, regardless of the lagged property in the LGN cells. However, intercortical inhibition without lagged LGN cells leads to a more symmetric (bidirectional) tuning, while additional lagged LGN cells lead to a dominant direction (unidirectional tuning).

**Figure 7:**
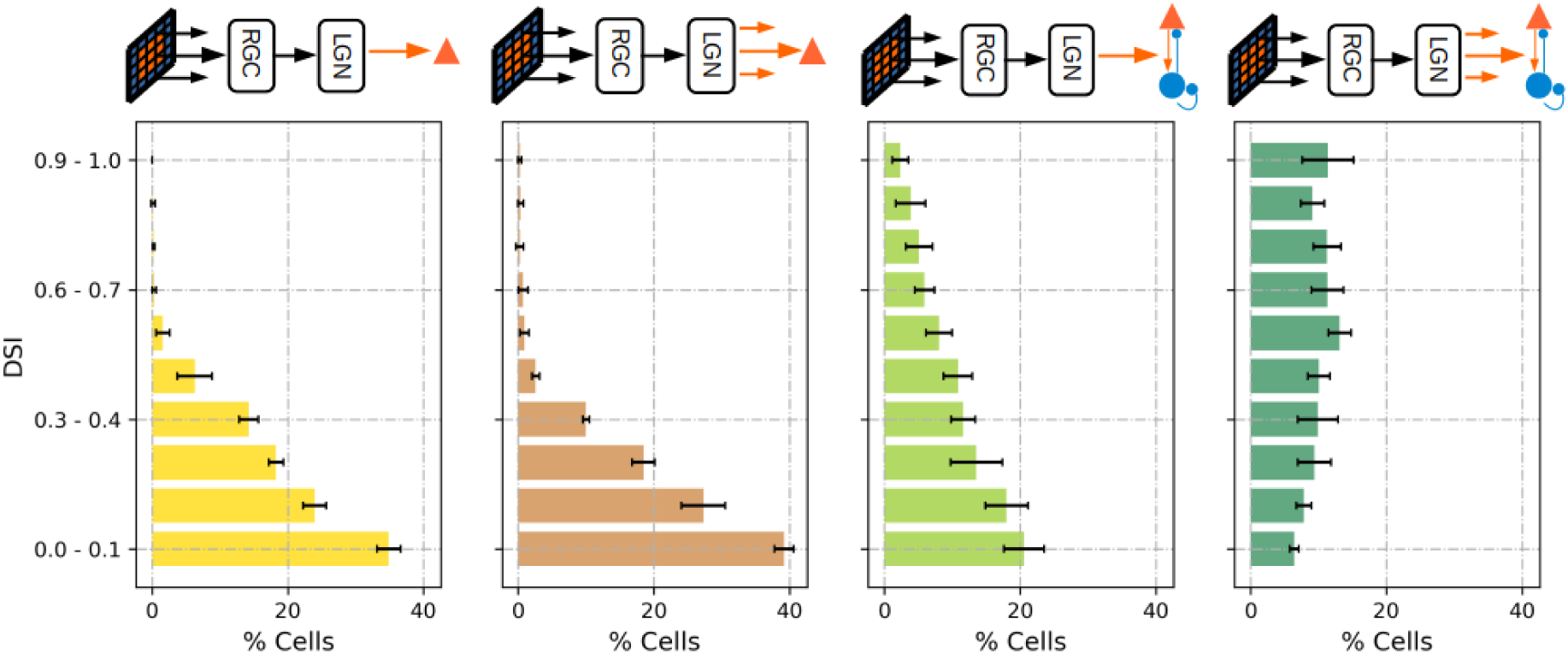
Inhibition and lagged LGN cells contribute to direction selectivity in V1. Percentage of simple cells in a certain range for the DSI. The length of each bar indicates the average percentage of cells over five network instances using different sets of pre-trained weights. The whiskers indicate the standard deviation over five network instances using a different set of pre-trained weights. Pictograms over the plots indicate lagged or non-lagged LGN cells (lagged: three orange arrows) or intercortical inhibition (blue lines) in the network. Without either intercortical inhibition or lagged LGN cells, the majority of simple cells showing no direction tuning (first plot on the left). With lagged LGN cells (but without inhibition), most simple cells are showing no direction tuning (second figure from left). With intercortical inhibition (but without lagged LGN cells), more cells show a weak to strong direction selectivity (third figure from left). With intercortical inhibition and lagged LGN cells, most of the cells show a strong unidirectional selectivity (fourth figure from left).

**Figure 8:**
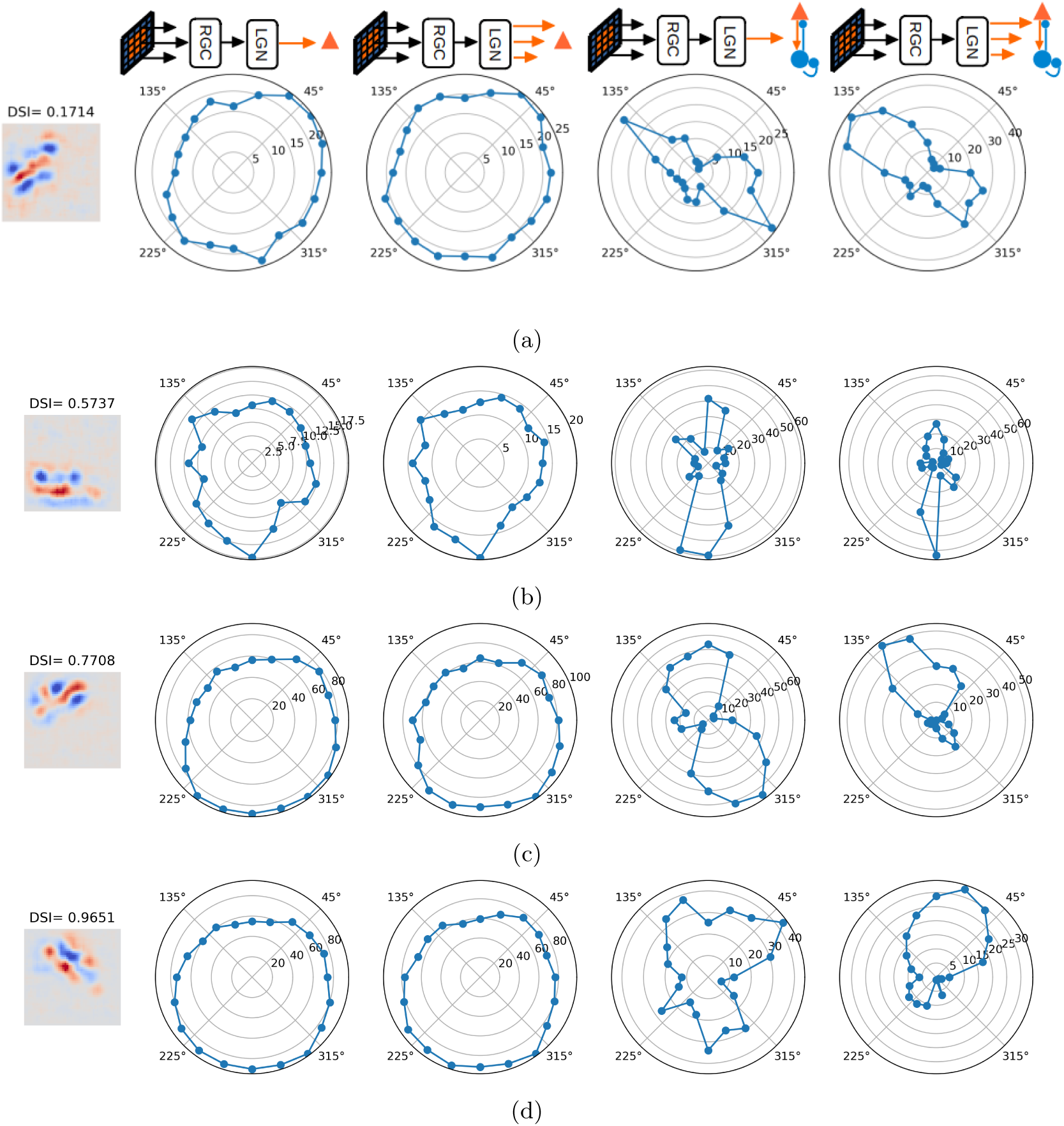
Response profiles to moving sinusoidal gratings of four cells with different levels of direction selectivity and different model configurations. Cells show a similar broadly tuned response profile in models without intercortical inhibition (first two columns). In models with inhibition (but without lagged LGN cells, third column), the response profile sharpens but is tuned to two opposing directions. In models with lagged LGN cells and inhibition (fourth column), the cells with a high DSI showed a clear preference for one of the two directions.

#### 3.2.1 Input offset leads to asynchronous inhibition

To better understand how inhibition and lagged cells support neuronal selectivity, we measured the temporal offset (Δ*T* ) between the excitatory and inhibitory input current and grouped the simple cells along their DSI value into 5 groups in the model with lagged cells and inhibition.

We observe that for cells with lower DSI values (between 0 and 0.2), the offset (Δ*T* ) for most of the cells is around -6 ms at the preferred stimulus direction (see **Fig. 9a**). A similar temporal offset is observable for the null direction, indicating, that the excitatory current arrives 6 ms before the inhibitory one for driving the neuron towards a spike, regardless of the stimulus direction. For cells with higher DSI, the temporal offset of the preferred direction changes to around -10 ms, indicating a longer temporal offset between the excitatory and inhibitory current, giving the cell enough time to spike before inhibition arrives, while on the null direction, the offset is close to 0 ms, indicating that the inhibitory and the excitatory currents arrive at nearly the same time to cancel each other out.

**Figure 9:**
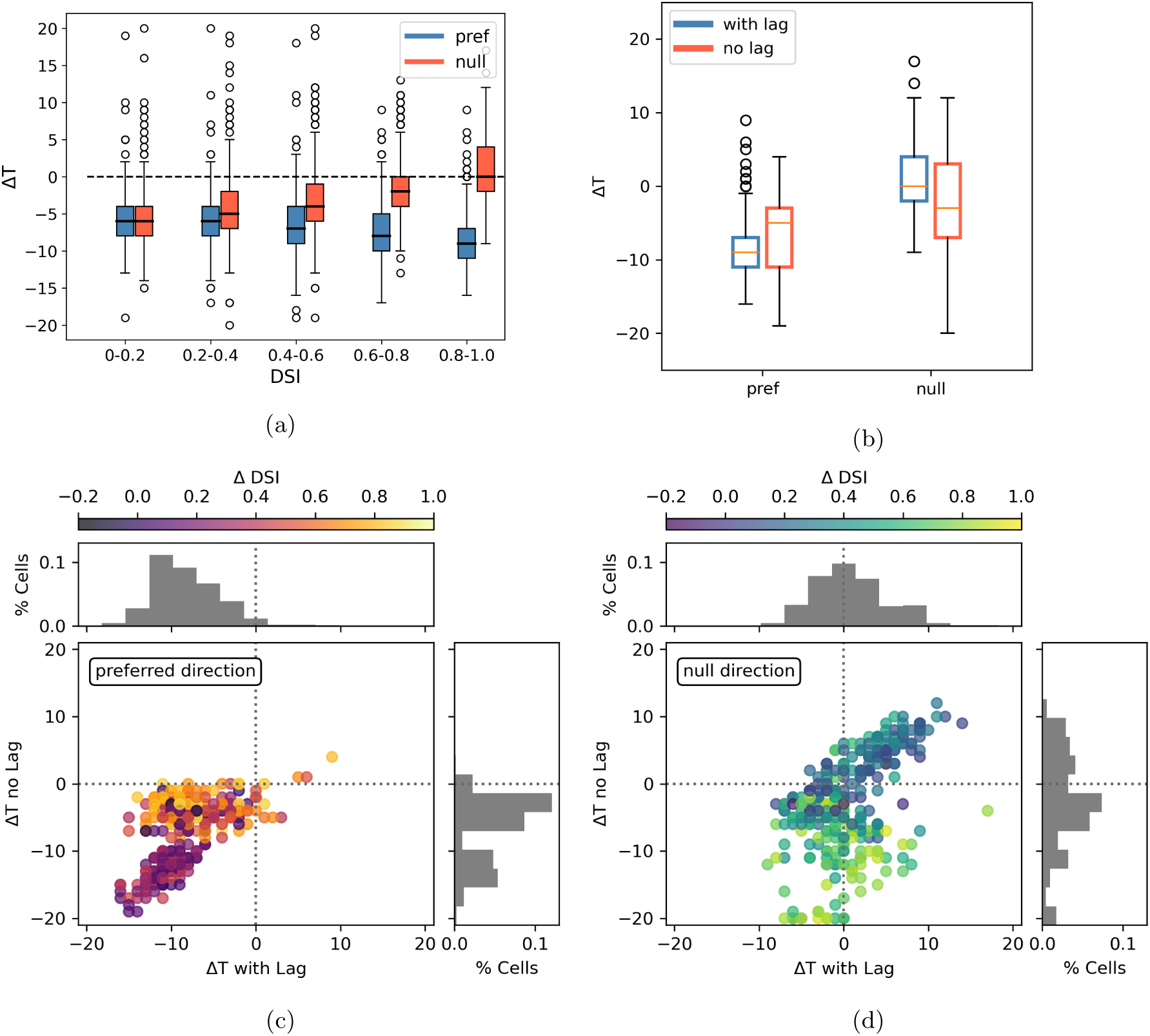
Temporal offset between excitatory and inhibitory input current is different for cells with a high DSI. Δ*T* marks the peak of the cross-correlation between the excitatory and inhibitory input currents to the simple cells. It indicates the offset between the excitatory and inhibitory current in ms. A negative Δ*T* means, that the inhibitory current arrives later than the excitatory one. (a) Temporal correlation between excitatory and inhibitory input currents as a function of DSI. For cells without a direction selectivity, the temporal offset for the preferred (blue) and non-preferred (orange) direction is equal (-6 ms). In cells with a high direction selectivity, the excitatory current arrives sooner than the inhibitory current at the preferred direction, otherwise, inhibition and excitation arrive at the same time. The black line indicates the median value. (b) Comparison of Δ*T* at the preferred direction and null direction for cells with a high direction selectivity with lagged LGN cells (blue) and without lagged LGN cells (red). With lagged LGN cells, the difference between the preferred and null direction is higher than without lagged LGN cells. (c) Δ*T* of cells that are highly direction selective with lagged LGN cells and without lagged LGN cells at the preferred direction. The color indicates the difference in the DSI between the model with lagged LGN cells and without. (d) Δ*T* of cells that are highly direction selective with lagged LGN cells and without lagged LGN cells at the null direction. The color indicates the difference in the DSI between the model with lagged LGN cells and without. Cells in (b), (c), and (d) are cells with a DSI *>* 0.8 in the model with intercortical inhibition and with lagged LGN cells.

To understand how the temporal offset and the direction selectivity are influenced by lagged LGN cells, we measured for simple cells with a high direction selectivity how the temporal offset and the direction selectivity changes (Δ DSI) if the response of lagged LGN is not delayed any more. We observe that for cells with a high Δ DSI value, inhibition and excitation arrive more synchronously at the preferred direction if no lagged LGN cells exist, while with lagged LGN cells inhibition arrives later than excitation (see **Fig. 9c**). In the null direction, we observe a reversed trend. Cells with a high Δ DSI value show a smaller temporal offset with lagged LGN cells, while without lagged LGN cells the absolute temporal offset gets higher, indicating a larger temporal gap between the excitatory and inhibitory input current (see **Fig. 9d**). Further, we observe that the temporal offset between the input currents without lagged LGN cells is more similar in both directions (see **Fig. 9b**). This suggests, that without lagged cells, both input currents in the preferred and the null direction arrive at a similar time and lead to a similar response in both directions. This symmetry is broken by a delay in the response of the lagged cells, leading to a later arrival of inhibition at the preferred direction and a synchronous arrival of both input currents at the null direction.

#### 3.2.2 Tuned inhibition improved direction selectivity

Due to the temporal offset, induced by the response delay of the lagged LGN cells, inhibition acts differently for the preferred direction compared to the null direction, leading to higher direction selectivity. However, as the pre-training on static natural scenes leads to structured connectivity between the excitatory and inhibitory population, as well as to tuned inhibitory cells, circuit motifs may also influence the direction selectivity. We calculated the DSI again for three model configurations: First, with a shuffled feedback connection from inhibition to excitation, excitatory neurons receive random feedback inhibition. Second, the feed-forward weights (from LGN and from the excitatory population) to the inhibitory populations are shuffled, leading to untuned inhibitory cells. Third, inhibitory feedback and feed-forward weights to the inhibitory population are shuffled. By comparing the resulting DSI values with the standard model, we observe small changes if only the inhibitory feedback weights are shuffled (see **Fig.10**). In contrast to that, shuffling the feedforward inhibition (leading to unselective inhibitory neurons), a strong decrease of selectivity for most of the cells with a high direction selectivity is observable. This suggests that the temporal difference between the excitatory and inhibitory input currents depends on the tuning of inhibitory and excitatory cells.

**Figure 10:**
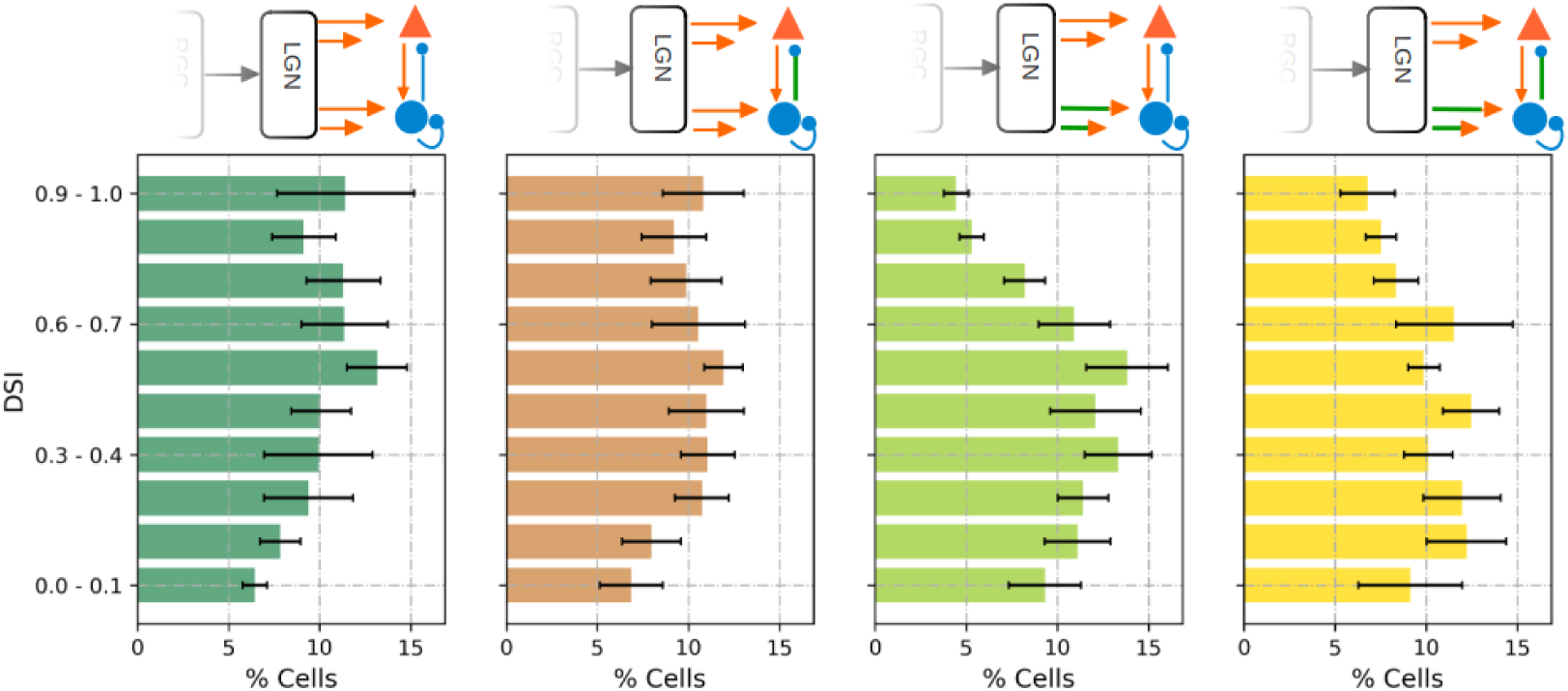
Tuned feed-forward inhibition contributes to direction selectivity. In models with tuned intercortical inhibition and with 50% lagged LGN cells with a delay of 50 ms, most of the cells show a high direction selectivity (first plot left). Untuned feedback inhibition does not change the DSI distribution (second plot from left). In contrast to that, untuned feed-forward inhibition leads to lower DSI values (third plot from left), and additional untuned feedback inhibition does not change the direction selectivity (fourth plot from left), indicating that tuned feed-forward inhibition mainly contributes to direction selectivity. The length of each bar indicates the average percentage of cells over five network instances. The whiskers indicate the standard deviation over five network instances.

To evaluate how the selectivity of excitatory and inhibitory neurons shape direction selectivity, we first calculated an approximated inhibitory current generated by each inhibitory neuron, which is sent to an excitatory neuron. For that, we multiplied the synaptic inhibitory feedback weight with the number of spikes in a time frame of 2 ms. Second, we calculated the temporal offset (respective to the Δ*T* as described above) between the excitatory input current of the excitatory cell and all approximated inhibitory currents. Third, we calculated the cosine similarity between the input synapses of all excitatory and inhibitory cells. Due to the fact, that the cell’s selectivity and its receptive fields are mainly controlled by the thalamocortical input from LGN, measuring the cosine similarity can be used to determine how similar the receptive fields of excitatory and inhibitory cells are. While a value closer to one indicates, that the inhibitory neurons have a similar orientation, and phase, and are in the same position in the input space, a negative similarity value indicates a receptive field with the same position and orientation but with a reverse phase. A similarity value around zero indicates mainly a receptive field with a different position at the input space (see **S1**). The cosine similarity is used to group the inhibitory cells in relation to the receptive field similarity of an excitatory neuron, to evaluate how the temporal offset between the excitatory and inhibitory input current is influenced by the receptive field similarity.

We observe that for highly direction-selective cells the input currents from inhibitory cells with a cosine around zero (e.g., inhibitory cells at a different position) have on average a temporal offset of Δ*T* = -4 ms, at the preferred direction and a Δ*T* = +5 ms at the null direction (see **Fig. 11**). This indicates that inhibitory neurons with a spatially distinct receptive field spiked later in the preferred direction and earlier in the null direction. In contrast to that, the inhibitory currents from cells with a spatially overlapping receptive field but with a similar orientation (what is true for inhibitory cells with a cosine around -0.8 and 0.8) arrive on average at a similar time as the excitatory input current regardless of the direction. For excitatory cells that do not show any level of direction selectivity (DSI *<* 0.2), the temporal offset is on average around zero, for both directions. These results indicate that the positional relation between excitatory and inhibitory cells also influences the direction selectivity.

**Figure 11:**
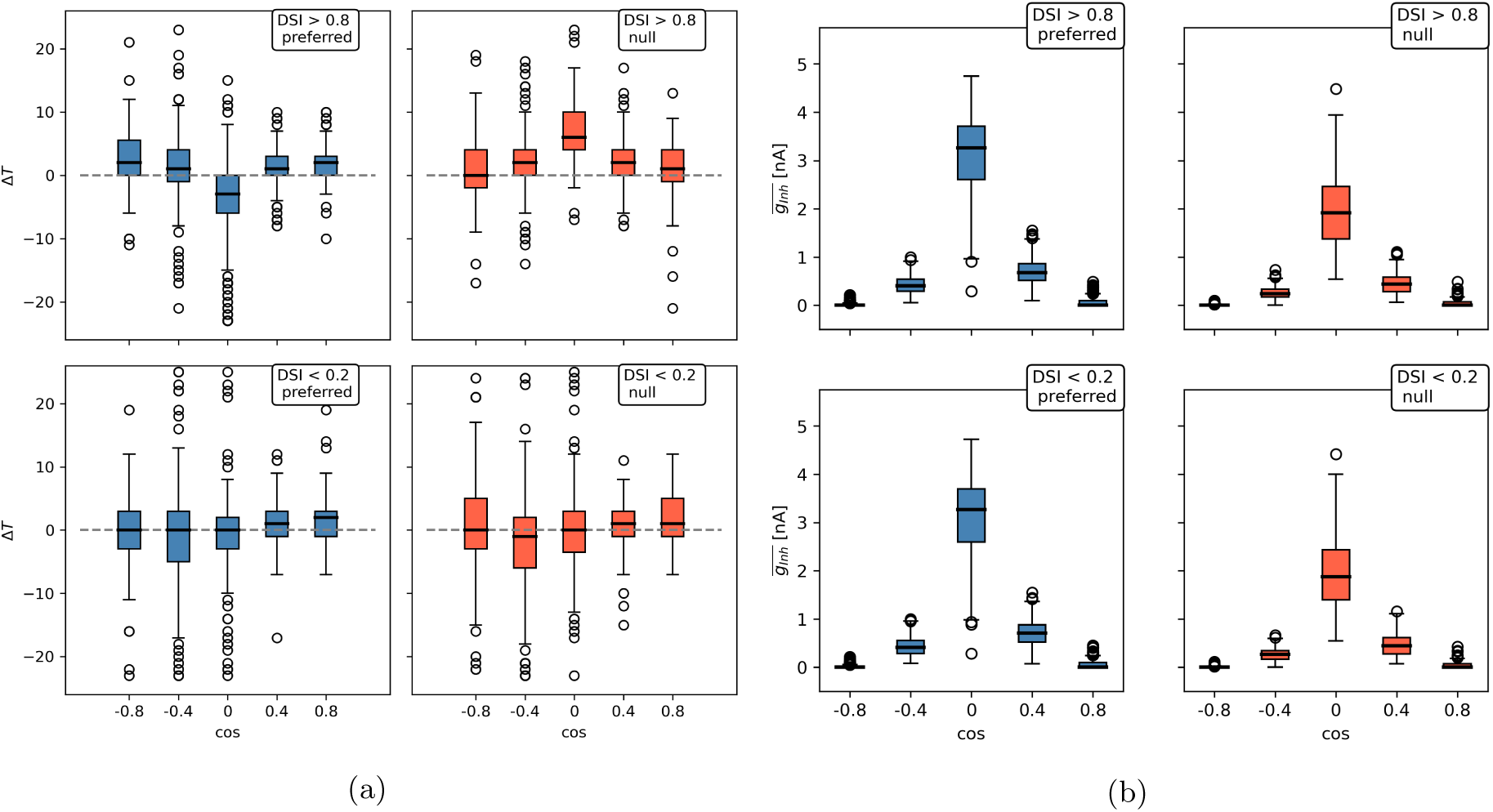
Tuned inhibition controls direction selectivity. (a) Cross-correlation between the excitatory input current and the approximated inhibitory input current as a function of the receptive field cosine. The Δ*T* marks the temporal shift in the inhibitory current that leads to the maximum correlation. For cells with a DSI *>* 0.8, the corresponding inhibitory current from cells with a cosine around zero shows a negative Δ*T*, indicating that this current arrives later than the excitatory current, while the inhibitory feedback from cells with a higher cosine value is nearly simultaneous. In contrast to that, in the null direction, the inhibitory current from cells with a cosine around zero arrives sooner than the excitatory input current. If the DSI *<* 0.2, the temporal difference between the cells is for both directions, preferred and null, similar. **(b)** Average approximated inhibitory current over the complete stimulus presentation as a function of the receptive field cosine. Independently of the DSI and the direction, most of the inhibitory current is generated by inhibitory cells with a receptive field having a different orientation and a spatial offset to the postsynaptic excitatory cell. In the average inhibitory current, there is no difference between cells with a high or low DSI. Top row, data for excitatory cells with a DSI *>* 0.8, and at the bottom row, data is presented for excitatory cells with a DSI *<* 0.2. On the left, data presented for the preferred orientation (in blue), and on the right data presented for the null direction (in red).

## 4 Discussion

Our study aimed at investigating how the tuning of intercortical inhibition and the temporal offset caused ny non-lagged and lagged LGN cells influence the emergence of direction selectivity by means of a neuro-computational model. Thus, we extended a biologically motivated spiking neural network of V1 layer 4 (Larisch et al., 2021) with populations of ON- and OFF-ganglion and LGN cells. By performing a systematic analysis of different network motifs, temporal, and neuronal dynamics, we identify how the characteristics of spatiotemporal receptive fields (STRFs) and direction selectivity of V1 simple cells emerges in the network.

### 4.1 Ganglion cell surround delay as the origin of spatiotemporal receptive fields

A large fraction of models implements spatiotemporal receptive fields (STRFs) by the combination of a spatial activation function and a non-linear bi-phasic function for the temporal behavior, either directly in the simple cell model (Adelson & Bergen, 1985; Yang et al., 2000; Lindeberg, 2011), in the model for LGN cells (Wimbauer, Wenisch, Miller, & van Hemmen, 1997; Hamada et al., 1997; Miyashita et al., 1997; Ursino et al., 2007; Okajima, 2014), or in the retinal ganglion cell model (McFarland et al., 2013; Mobarhan et al., 2018). Despite their success in modeling the functional components of STRFs and providing suitable models for further analysis (like for direction selectivity or the analysis of higher cortical areas), they do not explain how STRFs emerge through network circuit motifs.

Our model analysis explains how a delay in the surround field of the retinal ganglion cells enables the emergence of STRFs. Previous studies successfully modeled STRFs for retinal ganglion cells and LGN by extending the classical DoG receptive field to simulate a delay in the surround field (Enroth-Cugell et al., 1983; Dawis et al., 1984; Cai et al., 1997). While Enroth-Cugell et al. (1983) changed the DoG formula to simulate the functional implication of a delay in the frequency domain, Cai et al. (1997) multiplied the center and surround Gaussian each with a temporal function to implement the spatiotemporal processing. By adding a delay parameter solely to the temporal function of the surround field, Cai et al. (1997) showed better fits for the STRFs of LGN cells. Instead of an additional temporal function to model the temporal dynamics of the ganglion cell responses, we directly delayed the transmission of the surround inputs in the synapse objects of the ANNarchy simulator. Neuronal responses in all populations of our network are modeled with the biologically plausible adaptive exponential integrate-and-fire model, leading to STRFs for V1 simple cells (DeAngelis et al., 1995, 1993a) and also for cells in the LGN population (**Fig. S2**) as reported in cat (DeAngelis et al., 1995; Cai et al., 1997).

### 4.2 The origin of surround delay dynamic

To implement the initial STRF dynamic, we used a delay for the surround response relative to the center in the ganglion cell populations of 30 up to 50 ms. While this is in line with findings in rodents (Tokutake & Freed, 2008), a shorter delay of 6 ms to 8 ms was reported for primates (Benardete & Kaplan, 1997; Kilavik et al., 2003). However, a comparably long surround delay (20 ms to 40 ms) has been used to fit the spatiotemporal dynamics of bipolar cells in the primate retina (Y. J. Kim et al., 2022), which provide the excitatory input to ganglion cells. As the ganglion cell also receives inhibitory input from amacrine cells, we assume that the simplification in our ganglion cell model made longer delays necessary. The retina consists of different types of photoreceptors, bipolar, horizontal, amacrine, and ganglion cells (see Baden et al. (2020) and Grünert & Martin (2020) for a review), leading to a complex network with different circuits that influence the response of ganglion cells. Especially the role of inhibitory amacrine cells in this network is not fully understood, as they show a high morphological diversity in mice (Baden et al., 2020) and primate (Grünert & Martin, 2020) and sending feedback inhibition to bipolar cells, lateral inhibition to other amacrine cells and providing ganglion cells with feedforward inhibition (Diamond, 2017; Franke et al., 2017). Additionally, it has been suggested that ganglion cells send feedback information to amacrine cells via electrical synapses, building a recurrent circuit and manipulating inhibition send to bipolar and ganglion cells (Vlasiuk & Asari, 2021). Due to this, we assume that the spatiotemporal response profile of ganglion cells could be caused by this complex interaction, even with a shorter delay between center and surround. Different model studies of LGN cells support this assumption by demonstrating how excitatory and inhibitory feedback to LGN can influence the spatial and temporal profile (Yousif & Denham, 2007; Mobarhan et al., 2018).

### 4.3 Lagged LGN cells lead to more inseparable STRFs

A STRF is called separable if its space-time profile can be described by two separable functions for the space and time dimensions, otherwise, it is called inseparable (Adelson & Bergen, 1985; DeAngelis et al., 1993a; McLean et al., 1994; Ohzawa et al., 1996). It has been assumed previously, that temporal offset created by the combined response of non-lagged and lagged LGN cells creates the inseparability of STRFs (Saul & Humphrey, 1992; McLean et al., 1994). Studies in the mammalian visual cortex showed that STRFs of simple cells lay in a spectrum from clearly separable to strongly inseparable (DeAngelis et al., 1993b). We proposed here a new method to measure the grade of inseparability by calculating the difference between the temporal frequency amplitudes in the Fourier spectrum. With this temporal frequency amplitude difference (TFD), we quantify the numbers of simple cells with a specific level of inseparability in the corresponding STRF, where a value closer to zero indicates a separable STRF and a value closer to one indicates a more inseparable STRF. Our results show, that in the absence of lagged LGN cells, a majority of simple cells have a TFD *<* 0.2 and thus, a separable STRF. Adding a delay to the response of 50% of the lagged LGN cells results in an increase of the TFD value for most of the cells, accompanied with the emergence of more inseparable STRFs for these cells.

### 4.4 Inhibition together with lagged LGN cells leading to direction selectivity

Lagged cells have been found in the lateral geniculate nucleus of cats (Saul & Humphrey, 1990; Cai et al., 1997; Saul & Feidler, 2002; Vigeland et al., 2013), monkeys (Saul, 2008b), and mice (Piscopo et al., 2013). Different model studies showed how the combination of the shifted input through non-lagged and lagged LGN cells creates a temporal offset in the input to layer 4, being responsible for the emergence of direction selectivity, even without inhibition in layer 4 (Wimbauer, Wenisch, Miller, & van Hemmen, 1997; Wimbauer, Wenisch, van Hemmen, & Miller, 1997; Okajima, 2014; Lien & Scanziani, 2018).

It has been suggested, that the temporal offset in the input stream from LGN to V1 could not only be implemented through non-lagged and lagged LGN cells but also by transient and sustained LGN cells (Chizhov & Merkulyeva, 2020). While the existence of these cells is well known (Cleland et al., 1971; Marrocco, 1976; Saul & Feidler, 2002), their relevance for direction selectivity in the visual cortex remains unclear. It has been shown in mice’s visual cortex, that direction-selective cells receive input from LGN cells with a transient response profile and from cells with a sustained response profile (Lien & Scanziani, 2018). A potential role of these LGN cells for direction selectivity as been suggested (Chizhov & Merkulyeva, 2020) as well as neglected (Chariker et al., 2021; Freeman, 2021) by recent model studies. Besides those properties of LGN cells, researchers pointed out an effect of intracortical inhibition on direction selectivity in the visual systems of cats (Monier et al., 2003; T. Kim & Freeman, 2016), mice (Lee et al., 2012; Li et al., 2014; Rossi et al., 2020), ferrets (Wilson et al., 2018), and monkeys (Livingstone, 1998), even questioning the role of lagged LGN cells through the need for exact timing of lagged and non-lagged cells to achieve direction selectivity (Priebe & Ferster, 2005). Some model studies shown that direction selectivity could emerge through intracortical inhibition, without the need for lagged LGN cells (Wenisch et al., 2005; Freeman, 2021).

Our model simulations support the assumption that both, lagged LGN cells (or related mechanisms) and intracortical inhibition, has to be taken into account to explain direction selectivity. While previous models suggest that the thalamocortical input provides a basis direction selectivity, which is enhanced through intercortical inhibition (Ursino et al., 2007; Chizhov & Merkulyeva, 2020), our model simulations suggest an alternative explanation. Our results show, that simple cells show a clear orientation selectivity through intercortical inhibition, but no direction selectivity, if lagged LGN cells are missing, due to a similar arrival of excitatory feedforward and inhibitory feedback currents for both directions. Direction selectivity in our model increases if the temporal offset introduced by lagged LGN cells leads to a delayed response of either inhibitory cells or excitatory cells to break the synchronicity between both input currents at the preferred and null direction in favor of one of them.

### 4.5 Orientation and spatial offset as source of asymmetric inhibition

While our results show that the temporal offset between non-lagged and lagged LGN cells provides an advantage for one direction, we observed that direction selectivity also benefits from tuned feedforward inhibition. Inhibitory cells with different orientations and different spatial positions contribute by sending their inhibitory currents early at the null direction to cancel out the excitatory input current and only later at the preferred direction, improving direction selectivity. This temporal asymmetric inhibition has been found in the visual cortex of monkeys (Livingstone, 1998), cats (Priebe & Ferster, 2005), and mice (Li et al., 2014) and its spatial origin is in line with findings in mouse visual cortex (Rossi et al., 2020). Furthermore, various model studies investigated the influence of asymmetric inhibition on direction selectivity (Ursino et al., 2007) and even demonstrated how intercortical inhibition leads to direction selectivity without a temporal offset in the LGN feedforward path (Freeman, 2021).

By demonstrating that direction selectivity emerges through inhibitory cells with the same orientation and either the same or the opposite phase, Ursino et al. (2007) suggested that the phase of the inhibitory neuron in relation to the excitatory neuron is not important. Supporting this claim, our results show that the inhibitory neurons with the same orientation and the same phase as well as the opposite phase spike nearly simultaneously with the excitatory cell at the preferred and null direction, supporting the assumption that the inhibitory phase is irrelevant for direction selectivity. In a recent model study, Freeman (2021) demonstrated in a rate-coded orientation map model trained by a developmental procedure how direction selectivity emerges through intercortical inhibition alone, without a temporal offset in the thalamocortical path. In their model, direction-selective simple cells are located at the borders between two orientation preference maps. Due to this, inhibitory cells in the neighborhood of the respective simple cell are selective for different orientations. As a consequence, the averaged spatial receptive field of the inhibitory neurons shows a spatial offset from the receptive field of the simple cell. This leads to an inhomogeneous inhibitory input current in time, arriving later at the preferred direction and earlier at the non-preferred direction. Similar to the findings of Freeman (2021), we observe in our network that the accumulated input from inhibitory cells with a spatial offset and different orientations arrives late in the preferred direction and early on in the non-preferred direction. However, the model of Freeman (2021) did not contain any spatiotemporal dynamics in the simple cell population or a temporal offset in the thalamocortical feedforward path, leaving it open to how these circuit motifs would influence the direction selectivity. Beyond the study of Freeman (2021), our network demonstrated how direction selectivity is influenced by the combined effect of a temporal offset in the thalamocortical path and tuned intercortical inhibition.

### 4.6 Role of ganglion cell direction selectivity for simple cell selectivity

As direction-selective ganglion cells have been reported in the retina of primate (Y. J. Kim et al., 2022) and mice (Weng et al., 2005; Hillier et al., 2017), their role for direction selectivity of simple cells in the primary visual cortex remains unclear. While direction-selective cells in the retina of mice are frequent (Kay et al., 2011; Sabbah et al., 2017), only *≈* 1.5% of cells in the monkey retina are direction selective (Y. J. Kim et al., 2022), indicating only a small influence on the simple cell selectivity. Further, a recent study in the ferret visual pathway shows that LGN cells did not contribute to the experience-based development of direction selectivity in the cortex (Stacy et al., 2023), suggesting an intercortical mechanism for the V1 simple cell selectivity. Additionally, a study on mice retina demonstrated that disrupting the ganglion cell direction selectivity by blocking inhibition from starburst amacrine cells does reduce the tuning for the four cardinal directions in layer 2/3 of V1, but not the total amount of direction-selective cells and leads to a more even distribution between all directions (Hillier et al., 2017). This indicates a mechanism for direction selectivity in V1 based on intercortical dynamics rather than caused by the direction tuning in the retina alone. Due to this, we did not consider direction selective cells in the retina in our model. These cells may rater only shape the distribution of the preferred directions, and can not hold responsible for the basic emergence of direction selectivity in V1.

### 4.7 Predictive models of simple cell direction selectivity

As stated above, different models about the origin of direction selectivity have been published over the years, focusing on the role of lagged LGN cells and inseparable simple cell STRFs (Adelson & Bergen, 1985; Hamada et al., 1997; Miyashita et al., 1997), suggesting a basis selectivity through lagged LGN cells and a sharpening effect by intercortical inhibition (Ursino et al., 2007; Okajima, 2014; Chariker et al., 2021), or suggesting intercortical inhibition alone to be responsible for the emergence of direction selectivity (Wenisch et al., 2005; Freeman, 2021).

In other recent studies, models were designed to perform a specific task or to predict neuronal responses from physiological experiments.Rideaux & Welchman (2021) used a three-layer network (one V1 layer, one MT layer, and a readout layer) to show the emergence of direction-selective cells by optimizing the weights with the backpropagation algorithm to predict the velocity of an image sequence. To process information in time, they used a three-dimensional input (two space dimensions and one time dimension) to the network to process this information with three-dimensional filter kernels. Due to this, the emergence of a three-dimensional convolutional kernel can be assumed as space-time receptive fields in their V1 layer. A similar approach was used by Singer et al. (2023), who used four stacks of convolutional neural networks. They successively optimized the weights in each stack using the backpropagation algorithm to predict the activity of the previous stack in a given time window, and used grayscale video data as initial input. By using a three-dimensional convolutional kernel, they were able to demonstrate the emergence of different spatial and temporal dynamics, such as Gabor-like simple cell receptive fields, spatiotemporal receptive fields, direction-selective cells, and cells selective for pattern rather than for components, as found in MT (Movshon et al., 1985; H. X. Wang & Movshon, 2016).

A more detailed model of the visual system has been presented by E. Y. Wang et al. (2023), which used the backpropagation algorithm to optimize the predictions of neuronal responses in different areas of the mouse visual cortex, recorded on videos. They used different types of layers as LSTM layers or 3D-convolutional layers to capture information over time inside of their network. Besides matching experimental data recorded on video, they also demonstrated the correct prediction of neuronal response on new input data, such as direction selectivity on pink noise (E. Y. Wang et al., 2023).

While these approaches succeed in recreating different dynamics and phenomenons of the visual system, they lack in explaining the roles of different circuit motifs inside of the network, as these models did not elucidate the role of intercortical inhibition, nor the necessity to capture temporal information for direction selectivity in the networks.

### 4.8 Motion processing in higher cortical areas and future work

Since the spatiotemporal processing and direction selective response of V1 simple cells is the starting point for cortical motion processing, our model can serve as a starting point for models involving higher areas related to motion processing, such as MT or MST (Kravitz et al., 2011). It has been shown that simple cells in V1 and cells in MT respond differently to a moving plaid stimulus, which is created by adding two sinusoidal gratings with different orientations. While cells in MT respond to the direction of motion of the whole plaid pattern, simple cells respond to the individual sinusoidal components (Movshon et al., 1985). In contrast to that, recent studies have reported the existence of cells responding to the plaid pattern in V1 of monkeys (Guo et al., 2004) and rats (Matteucci et al., 2023). Furthermore, a study in mouse visual cortex showed that the majority of cells in V1 are pattern selective rather than component selective (Palagina et al., 2017). Since several model studies have shown that feedback from MT to V1 influences V1 responses, and can create a feedback loop that improves flow and motion estimation in MT (Bayerl & Neumann, 2004; Raudies et al., 2011; Löhr et al., 2019), we assume feedback as a possible mechanism to explain the occurrence of pattern and component selective cells in V1. To evaluate this hypothesis, our network could be extended to include a population of MT cells and feedback synapses.

### 4.9 Conclusion

In this study, we presented a systematic analysis of different network motifs and neuronal dynamics in a spiking neural network. We demonstrated how these motifs incorporate with each other to shape the characteristics of spatiotemporal receptive fields (STRFs) and to enable direction selectivity for V1 simple cells (see **Fig. 12** for a summary). We demonstrated how a delay in the surrounding field of ganglion cells solely leads to the emergence of separable STRFs for cells in the LGN, as well as for V1 simple cells. Further, we showed how the addition of a response delay to 50% of the LGN cells leads to the emergence of inseparable STRFs in the V1 simple cell population. We systematically analyzed the influence of the temporal offset in the thalamocortical input path and the influence of tuned and non-tuned intercortical inhibition on direction selectivity. Direction selectivity emerges due to two mechanisms in our network: First, tuned lateral inhibition is mainly influenced by the preferred orientation and spatial receptive field offset, leading to a weak direction selectivity of most cells. Together with the second mechanism, the temporal offset caused by non-lagged and lagged LGN cells gives an additional temporal advantage and together with inhibition, tunes the response in the preferred and null direction.

**Figure 12:**
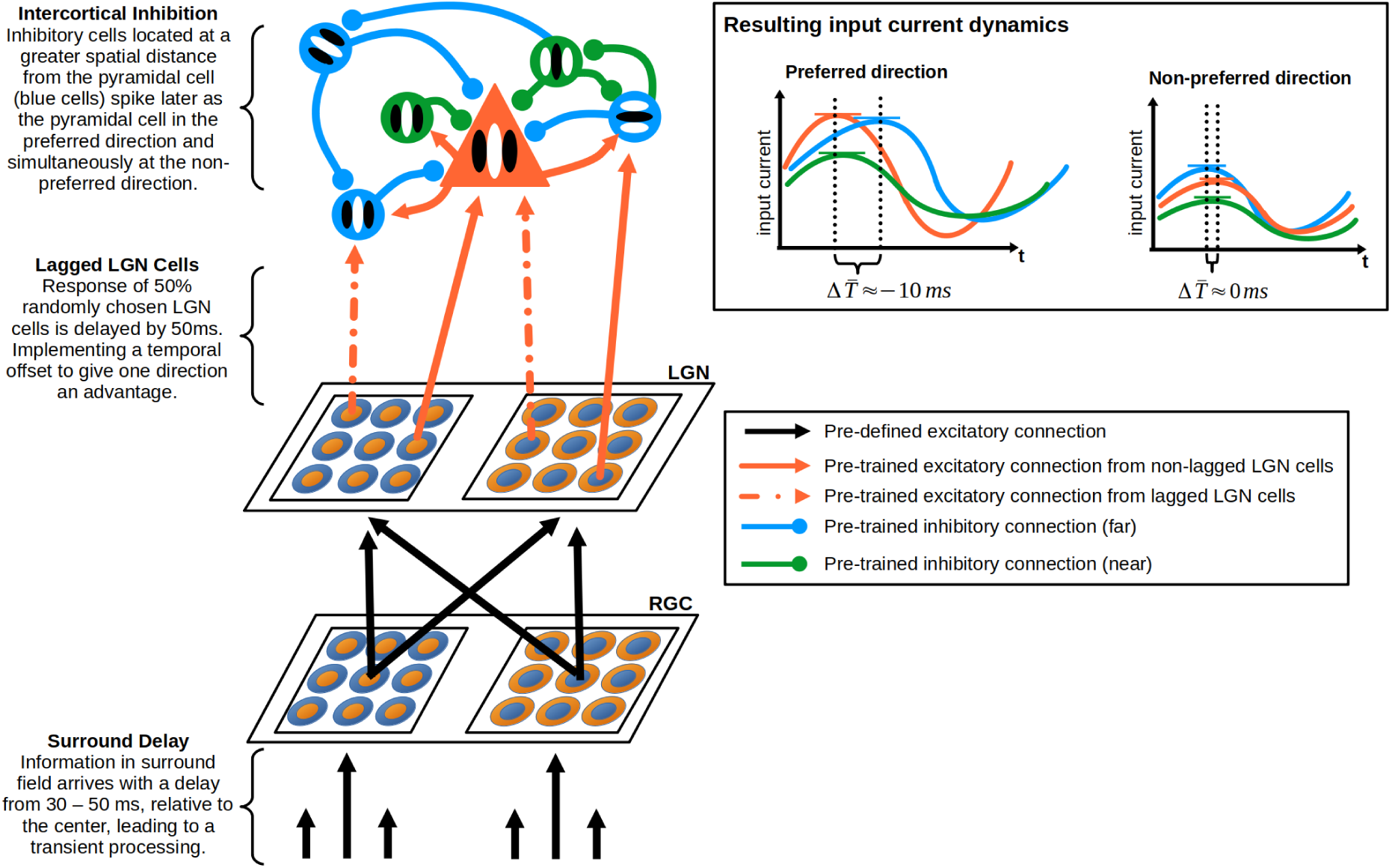
Network motifs and their role for direction selectivity. Due to the delays in the surround field of the retinal ganglion cells, spatial information of the centre and surround are decoupled in time, leading to transient processing in the next layer. A temporal offset between lagged and non-lagged LGN cells leads to a delayed response and dependent on the thalamocortical connectivity will give one direction a temporal advantage. Tuned intercortical inhibition increases direction selectivity, as inhibitory cells with a higher spatial offset of their receptive field and a different orientation spike differently at the preferred and non-preferred direction. Together, these network motifs influence the temporal dynamics of excitatory and inhibitory input currents to enable direction selectivity.

## Acknowledgment

This research has been partly funded by Horizon Europe Framework Programme, Awareness inside: EIC Pathfinder Challenges 2021, Grant Agreement 101071178 (CAVAA).

## Author Contributions

Conceptualization Reńe Larisch

Formal Analysis Reńe Larisch

Funding Acquisition Fred H. Hamker

Investigation Reńe Larisch

Methodology Reńe Larisch

Resources Fred H. Hamker

Software Reńe Larisch

Validation Reńe Larisch

Visualization Reńe Larisch

Writing - Original Draft Preparation Reńe Larisch

Writing - Review & Editing Fred H. Hamker, Reńe Larisch

